# FISHtoFigure: An easy-to-use tool for rapid, multi-target partitioning and analysis of sub-cellular mRNA transcripts in smFISH data

**DOI:** 10.1101/2023.06.28.546871

**Authors:** Calum Bentley-Abbot, Rhiannon Heslop, Chiara Pirillo, Matthew C. Sinton, Praveena Chandrasegaran, Gail McConnell, Ed Roberts, Edward Hutchinson, Annette MacLeod

## Abstract

Single molecule fluorescence *in situ* hybridisation (smFISH) has become a valuable tool to investigate the mRNA expression of single cells. However, it requires a considerable amount of bioinformatic expertise to use currently available open-source analytical software packages to extract and analyse quantitative data about transcript expression. Here, we present FISHtoFigure, a new software tool developed specifically for the analysis of mRNA abundance and co-expression in QuPath-quantified, multi-labelled smFISH data. FISHtoFigure facilitates the automated spatial analysis of transcripts of interest, allowing users to analyse populations of cells positive for specific combinations of mRNA targets without the need for bioinformatics expertise. As a proof of concept and to demonstrate the capabilities of this new research tool, we have validated FISHtoFigure in multiple biological systems. We used FISHtoFigure to identify an upregulation of T-cells in the spleens of mice infected with influenza A virus, before analysing more complex data showing crosstalk between microglia and regulatory B-cells in the brains of mice infected with *Trypanosoma brucei brucei*. These analyses demonstrate the ease of analysing cell expression profiles using FISHtoFigure and the value of this new tool in the field of smFISH data analysis.

## Introduction

Single molecule fluorescence *in situ* hybridisation (smFISH) technologies such as RNAScope enable the visualisation of single mRNA molecules within single cells. mRNA transcripts are detected by fluorescence microscopy, with each transcript appearing as a single ‘transcriptional spot’ (1). Quantification of these signals enables the analysis of transcriptional activity at the single cell level within the spatial context of tissues (2). However, the large microscopy datasets produced by smFISH experiments currently require custom code in order to conduct in-depth transcriptomic analyses. QuPath is a purpose-built platform for the analysis of large images such as those acquired during smFISH experiments, and is recommended by ACDBio-Techne, the developer of the RNAScope platform (https://acdbio.com/qupath-rna-ish-analysis), for image analysis (3). QuPath has specific in-built tools for cell segmentation and fluorescent spot detection, which can be used to quantify transcriptional spots. Furthermore, the software incorporates a batch processing feature which facilitates automated analysis of data from multiple images (3). Following quantification, QuPath can plot quantified data, such as transcripts per cell, as a histogram (3). However, users wishing to conduct more complex analyses, such as differential expression analysis or co-expression analysis, must develop custom pipelines to parse raw QuPath output data, thus restricting such analysis to users with extensive bioinformatics experience.

Here, we present FISHtoFigure, a standalone, open-source software tool for the in-depth analysis of transcript abundance in QuPath-quantified smFISH data by users with all levels of bioinformatic experience. FISHtoFigure can concatenate the batch processed data from QuPath, enabling the analysis of large, multi-image datasets. Notably, FISHtoFigure allows users to conduct transcript abundance analysis for cells with specific, multi-transcript expression profiles. Additionally, FISHtoFigure enables users to conduct differential expression analysis between datasets, facilitating the targeted study of differential expression in specific cell types and populations. Thus, FISHtoFigure provides a means for all users to examine mRNA expression of multiple transcripts without the need for custom analysis pipelines.

Here, we demonstrate the use of FISHtoFigure in two biological scenarios. First, we used FISHtoFigure to analyse T-cell and B-cell populations in the spleens of influenza A virus (IAV) infected mice, hereafter referred to as the spleen dataset. Second, we demonstrate the capabilities of FISHtoFigure for the analysis of high-plex smFISH data collected from highly ramified, non-round cell types, using a dataset obtained in a recent experiment by our group investigating microglia in the brains of *Trypanosoma brucei brucei* infected mice, hereafter referred to as the brain dataset (4).

## Materials and Methods

### Specifications, Usage, and Data Handling

FISHtoFigure is a Python-based analytical software tool, designed to quantify cell expression profiles within smFISH data. Expression profile analysis is conducted in FISHtoFigure using the Pandas library (5). A two-branched strategy is used to isolate cellular and subcellular data into two new datasets (**Figure 1A**). Graphical outputs are generated using a combination of the Matplotlib and Seaborn Python libraries (6,7). In addition to the graphical outputs, data from FISHtoFigure analysis are stored in CSV format for downstream statistical analysis. The statistical tests in this paper were performed with GraphPad PRISM.

**Figure 1:**
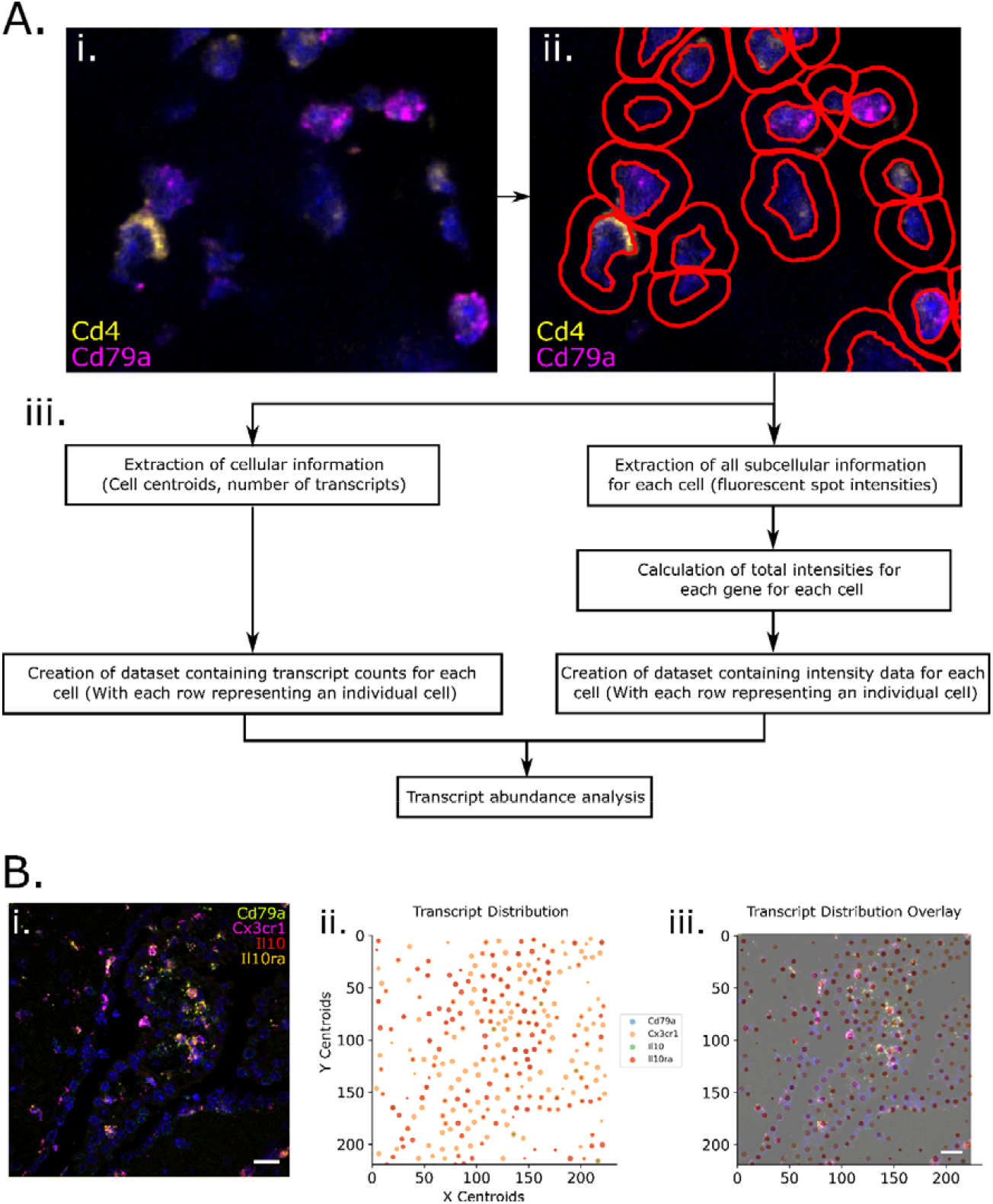
FISHtoFigure pipeline. **A**: [**i**] An smFISH image from the spleen dataset captured via confocal microscopy (Zeiss LSM 880). [**ii**] QuPath’s “Cell Detection” function was used to identify cell boundaries (shown in red). Cell nuclei identification is based on fluorescence above background in the channel associated with the nuclear stain (DAPI). [**iii**] An overview of FISHtoFigure processing of QuPath output data to generate transcript abundance outputs. **B**: [**i**] An smFISH image from the brain dataset (scale bar = 20µm), captured by confocal microscopy (Zeiss LSM 710) and [**ii**] processed using FISHtoFigure’s “Plot transcript distribution” function, where points represent cells and are sized based on the number of transcripts being expressed by that cell. [**iii**] An overlay of the captured smFISH image with the plot produced by FISHtoFigure demonstrates the accuracy of the pipeline.

### Sample Collection

All spleen samples were collected from 9 week old, male C57BL/6 mice. A mouse was infected with the IAV A/Puerto Rico/8/34 (PR8; H1N1) and samples were harvested 6 days post infection. Samples from an uninfected male C57BL/6 mouse were harvested to act as naïve controls. All brain samples were collected from 6 – 8-week-old, female C57BL/6 mice. Two mice were infected with *T. b. brucei* Antat 1.1E (4). Samples were harvested at 45 days post infection. Samples from two uninfected female C57BL/6 mice were harvested to act as naïve controls.

All samples were fixed in 4% paraformaldehyde (PFA) at room temperature for 24 hours and embedded in paraffin. From paraffin blocks, sections were cut on a microtome (Thermo Scientific) and mounted on glass slides for histology. All animal work was carried out in line with the EU Directive 2010/63/eu and Animal (Scientific Procedures) Act 1986, under project licences P72BA642F (Spleen samples) and PC8C3B25C (Brain samples), and was approved by the University of Glasgow Animal Welfare and Ethics Review Board.

### RNAScope data collection

Commercial RNAScope control slides containing mouse NIH 3T3 cells (Advanced Cell Diagnostics, US) were used as a positive control sample for RNAScope for all samples.

RNAScope was used to visualise *Cd79a* and *Cd4* transcripts in the spleens of naïve and IAV infected mice, and *Cd79a, Cx3cr1, Il10* and *Il10ra* transcripts in the brain of nailve and *T. b. brucei* infected mice. Fresh probe mixes containing the RNAScope probes were prepared for each experiment (**Table 1**). A single probe per channel (C1-C4) was included in each experiment. RNAScope 4-plex positive controls (for *Polr2a, Ppib*, and *Ubc*) and negative controls (for the *Bacillus subtilis* bacterial *Dapb* gene) were also included (probe details are listed at https://acdbio.com/control-slides-and-control-probes-rnascope). Slides were imaged by confocal microscopy (Zeiss LSM 880, 63x objective for the spleen samples; Zeiss LSM 710, 63x objective for the brain samples) within 72 hours of staining.

**Table 1.** RNAScope targets and detection specifications.

| Target | Channel | Detection dye | Dye dilution | Detection Channel |
| --- | --- | --- | --- | --- |
| ***Pol2ra*** | C1 | Opal 520 | 1:1500 | FITC |
| ***Ppib*** | C2 | Opal 570 | 1:1500 | Cy3 |
| ***Ubc*** | C3 | Opal 690 | 1:1500 | Cy5.5 |
| ***Dapb*** | C1, C2, C3 | Opal 520/570/690 | 1:1500 | FITC, Cy3, Cy5.5 |
| ***Cd79a*** | C1 | Opal 520 | 1:1500 | FITC |
| ***Cd4*** | C4 | Opal 650 | 1:1500 | PE/Cy5 |
| ***Cxc3r1*** | C2 | Opal 650 | 1:1500 | PE/Cy5 |
| ***Il10*** | C3 | Opal 570 | 1:750 | Cy3 |
| ***Il10ra*** | C4 | Opal 540 | 1:1500 | FITC/Cy3 |

### QuPath Image analysis

Once imaged, QuPath 0.3.1 Software was used to quantify the number of transcripts for each target probe (3). Negative control images were generated by probing spleen and brain tissue sections with the RNAScope 3-plex negative control probes. Fluorescence measurements for each detection channel in the negative control images were subtracted from final experimental images to determine fluorescence thresholds. Using in-built QuPath annotation tools, one large region of interest (ROI) was specified on each image such that the whole image was encompassed in a single annotation. The “Cell Detection” function was used to determine the number and position of cells in each ROI based on the DAPI nuclear stain (under the assumption that one nucleus represented one cell), and the ‘Subcellular Detection’ function was used to calculate the number of transcripts for each target. QuPath output data were then used as input data for FISHtoFigure. The analysis workflow was scripted to enable batch processing of all images within each dataset.

## Results

We designed FISHtoFigure to facilitate the conversion of QuPath-quantified image data into transcript abundance analytics. We designed a simple graphical user interface and packaged the FISHtoFigure software as a standalone executable program, enabling analysis to be conducted with no interaction with the raw data or underlying Python code. Below we outline the steps involved in analysing smFISH data using FISHtoFigure, along with examples of analysis outcomes.

### Step 1: Data Harvesting and Validation of Quantified smFISH Data

First, cellular boundaries and mRNA transcripts were identified using QuPath. QuPath output data were then processed using FISHtoFigure to produce differential transcript abundance analytics for different cell types or expression profiles (3). An overview of the FISHtoFigure pipeline is given in **Figure 1A** and an example of a typical image for processing is given in **Figure 1Bi**.

As experiments usually require numerous individual images, we created a dedicated pre-processing tool to concatenate individual QuPath-quantified image datasets into a single file comprising data from any number of smFISH images, which can then be analysed by the main FISHtoFigure program. Due to the volume of information captured during imaging, the resulting quantified files are large and include metrics not relevant for transcript expression analysis (e.g. morphometric data, such as, cell area, nucleus and cytoplasm morphology, etc.). The desired information, i.e., the number of transcripts per cell and fluorescent intensity data, which comprise only a small portion of the quantified data, were extracted by FISHtoFigure from QuPath-quantified smFISH data files and assigned to the cells from which they originate. Metrics were then calculated for each cell, i.e. the number of transcripts and total fluorescent intensities for each mRNA target. In addition to transcriptome information, cell location information is extracted in the form of the cell centroid (based on nuclear staining identified using the “Cell Detection” function in QuPath). These data are then processed by FISHtoFigure using the “Plot Transcript Distribution” feature, which produces a scatter plot of points representing cell centroids, with points sized by number of mRNA transcripts within the cell and coloured by gene (**Figure 1Bii**). This allows users to visualise quantified data in a format analogous to the original smFISH image (**Figure 1Bi**) and, by overlaying this visualised data with the original smFISH image, directly validate the accuracy of data extraction by FISHtoFigure (**Figure 1Biii**).

### Step 2: Differential target abundance analysis from smFISH data using the FISHtoFigure package

Following data extraction and assignment of transcript information to cells, differential transcript abundance analysis can be conducted using FISHtoFigure’s “Transcript abundance analysis” feature.

Using our spleen dataset, we investigated T-cell and B-cell populations in the spleens of mice, either uninfected or 6 days after infection with influenza A virus. These cells are highly abundant in spleen tissue and have a classically “round” cellular morphology. Their morphology enabled easy identification of cell boundaries in QuPath, and thus generated a straightforward dataset for software validation. Spleen sections from naïve and infected mice were stained using DAPI to identify cell nuclei and probed for *Cd4* and *Cd79a* mRNA transcripts, enabling us to identify helper T-cells and B-cells, respectively (8,9). This analysis revealed a statistically significant upregulation of *Cd4* expression during infection (p<0.01, Mann-Whitney test; **Figure 2A**), while no statistically significant difference in *Cd79a* expression was observed. In addition to graphical outputs, FISHtoFigure analysis is saved in CSV format for further downstream analysis. Here, statistical analysis was performed on the FISHtoFigure output data using GraphPad PRISM.

**Figure 2:**
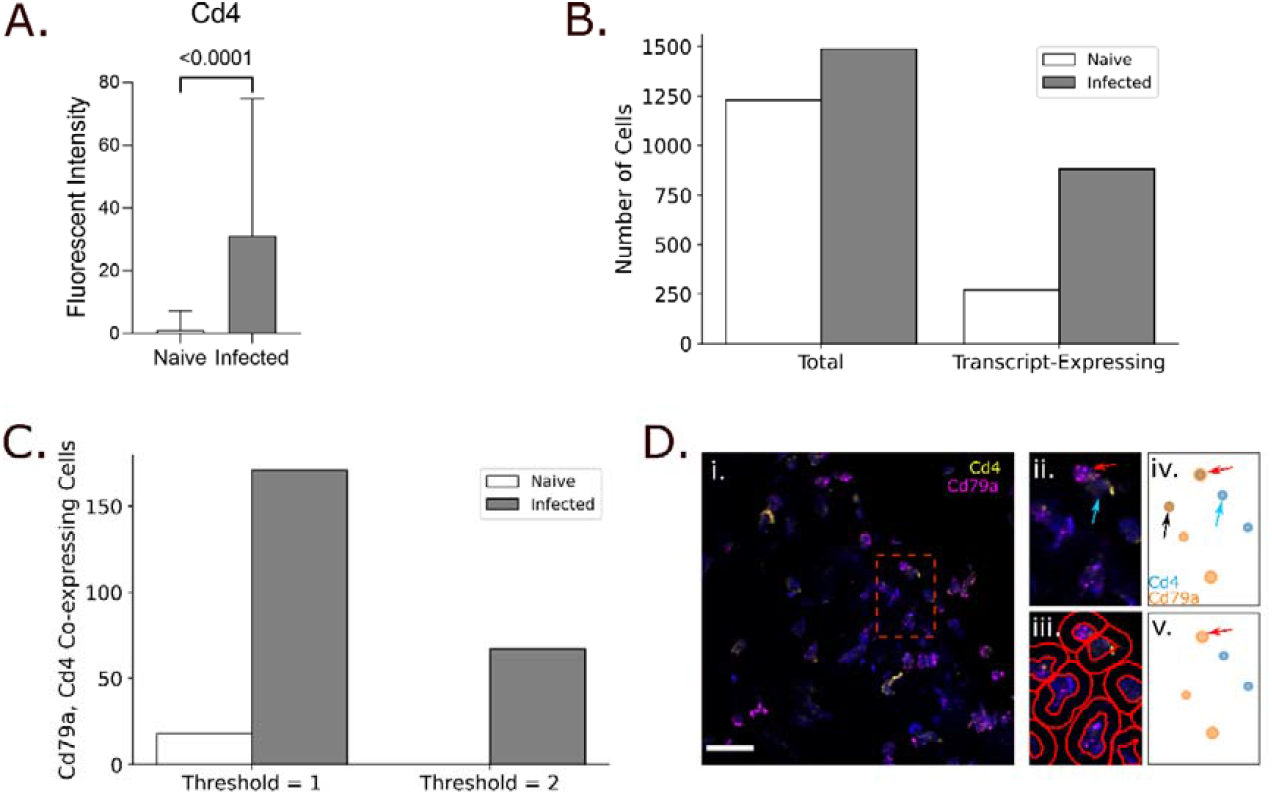
Analysis of spleen samples from naïve and influenza A virus infected mice. **A**: FISHtoFigure quantification of *Cd4* expression in the naïve and infected spleen datasets, significantly upregulated during infection (Mann-Whitney test). **B**: Total number of cells and the number of cells expressing at least one transcript within the naïve and infected spleen datasets. **C**: Number of cells co-expressing *Cd79a* and *Cd4*, with threshold set to 1 or 2 transcripts. **D**: [**i**] An smFISH image from a naïve spleen (scale bar = 20µm), captured by confocal microscopy (Zeiss LSM 880). [**ii**] A zoomed view of the region shown in the red square in [**i**] shows a B-cell (red arrow) and T-cell (blue arrow) in close proximity. [**iii**] Cell boundaries identified using QuPath. [**iv**] FISHtoFigure’s “Plot Transcript Distribution” feature with a threshold of 1 transcript per cell, [**v**] with a threshold of 2 transcripts per cell. Setting a threshold of 2 transcripts per target per cell results in the B-cell being correctly categorised (red arrow) – note the removal of the ambiguous *Cd79a*^*+*^ *Cd4*^*+*^ cell expressing both transcripts as they are below threshold levels (black arrow in [**iv**]).

We expanded the analysis capabilities of FISHtoFigure by adding the “Multi-target transcript abundance” feature, enabling the identification and quantification of cell types with multiplex transcriptomic profiles. This feature can be used to identify cells expressing any combination of mRNA transcripts. Here, we used this feature to validate the cell type quantification of our pipeline. Cd4 and Cd79a are well established markers for helper T-cells and B-cells respectively (8,9). Spleen resident B-cells do not express *Cd4*, and T-cells do not express Cd79a. Therefore we used the double-positive *Cd4*^+^ *Cd79a*^+^ cell population as a metric for mis-categorisation of cells by FISHtoFigure. The naïve dataset comprised a total of 1229 cells of which 273 contained transcripts of *Cd4* or *Cd79a* (**Figure 2B**). A total of 18 cells were labelled as *Cd4*^+^ *Cd79a*^+^, representing approximately 1.5% of all cells and 6.6% of transcript-expressing cells (**Figure 2C**). The infected dataset comprised a total of 1487 cells of which 882 contained transcripts (**Figure 2B**). The infected dataset showed a higher presumed mis-categorisation rate, with 171 cells (11.5% of all cells and 19.4% of transcript-expressing cells) labelled as *Cd4*^+^ *Cd79a*^*+*^ (**Figure 2C**).

Upon closer inspection of the quantified data, many of the apparently *Cd4*^+^ *Cd79a*^+^cells contained a majority of transcripts from one gene, suggesting that mis-categorisation typically resulted from a small number of transcripts from the other gene. This could be plausibly explained if incorrect boundary approximations caused a small proportion of transcripts to be mis-allocated between highly localised cells. For example, a B-cell in close proximity to T-cell might appear to contain a single *Cd4* transcript due to cell boundary approximation (**Figure 2D**). In such cases, it is reasonable to assume the cell identity based on the majority transcript. To address this, we introduced a thresholding feature so that users can define the minimum number of transcripts from each mRNA target required for cells to be included in analysis. By setting this threshold at 2 transcripts from each mRNA, the population of *Cd4*^+^ *Cd79a*^+^ cells was eliminated in the naïve dataset and substantially reduced (67 cells, representing 4.5% of all cells and 7.5% of transcript-expressing cells) in the infected dataset (**Figure 2C**). This was consistent with the model that *Cd4*^+^ *Cd79a*^+^ cells were artefacts, and showed that thresholding allowed this source of error to be controlled.

Having demonstrated that FISHtoFigure can quantify cell types based on mRNA expression profiles, we progressed to a more challenging system containing cells with less regular boundaries. To do this we examined sections of mouse brains, which contain highly ramified cell types, using data from a study exploring the interactions between regulatory B-cells (Bregs) and microglia during infection with *T. b. brucei* (4). The brain dataset comprised 17 images captured from brain sections of infected mice and 9 captured from uninfected (naïve) controls. Brain sections were stained using DAPI and probed for *Cd79a* (a B-cell marker), *Cx3cr1* (a microglia marker), *Il10* (an anti-inflammatory cytokine hypothesised to be involved in Breg–microglia interactions), and *Il10ra* (the receptor for *Il10*) (9,10). These images were quantified in QuPath and concatenated into two datasets comprising naïve control data and infected data.

Cx3cr1 is a well-established microglia marker (10). B-cells do not express *Cx3cr1* and microglia do not express *Cd79a*. Similarly to the spleen dataset, in order to examine to what extent the thresholding function could improve cell type quantification in data containing ramified cells, presumptively mis-categorised *Cd79a*^*+*^ *Cx3cr1*^*+*^ cells were quantified. The naïve dataset contained 1631 cells, 914 of which contained transcripts. 30 cells were labelled *Cd79a*^*+*^ *Cx3cr1*^+^ double-positive (1.8% of all cells, 3.3% of transcript-expressing cells). The infected dataset contained 3907 cells, of which 3332 contained transcripts, 392 were labelled as *Cd79a*^*+*^ *Cx3cr1*^+^ double-positive (10% of all cells, 11.7% of transcript-expressing cells; **Figure 3A**). Applying a threshold of 2 transcripts per mRNA per cell reduced the number of *Cd79a*^*+*^ *Cx3cr1*^*+*^ double-positive cells to 4 in the naïve dataset (0.2% of all cells, 0.4% of transcript-expressing cells), and 76 in the infected dataset (1.9% of all cells, 2.3% of transcript-expressing cells; **Figure 3A**). This demonstrated that applying thresholds for transcript abundance could allow accurate allocation of transcripts to cells even for cells with complex and irregular boundaries.

**Figure 3:**
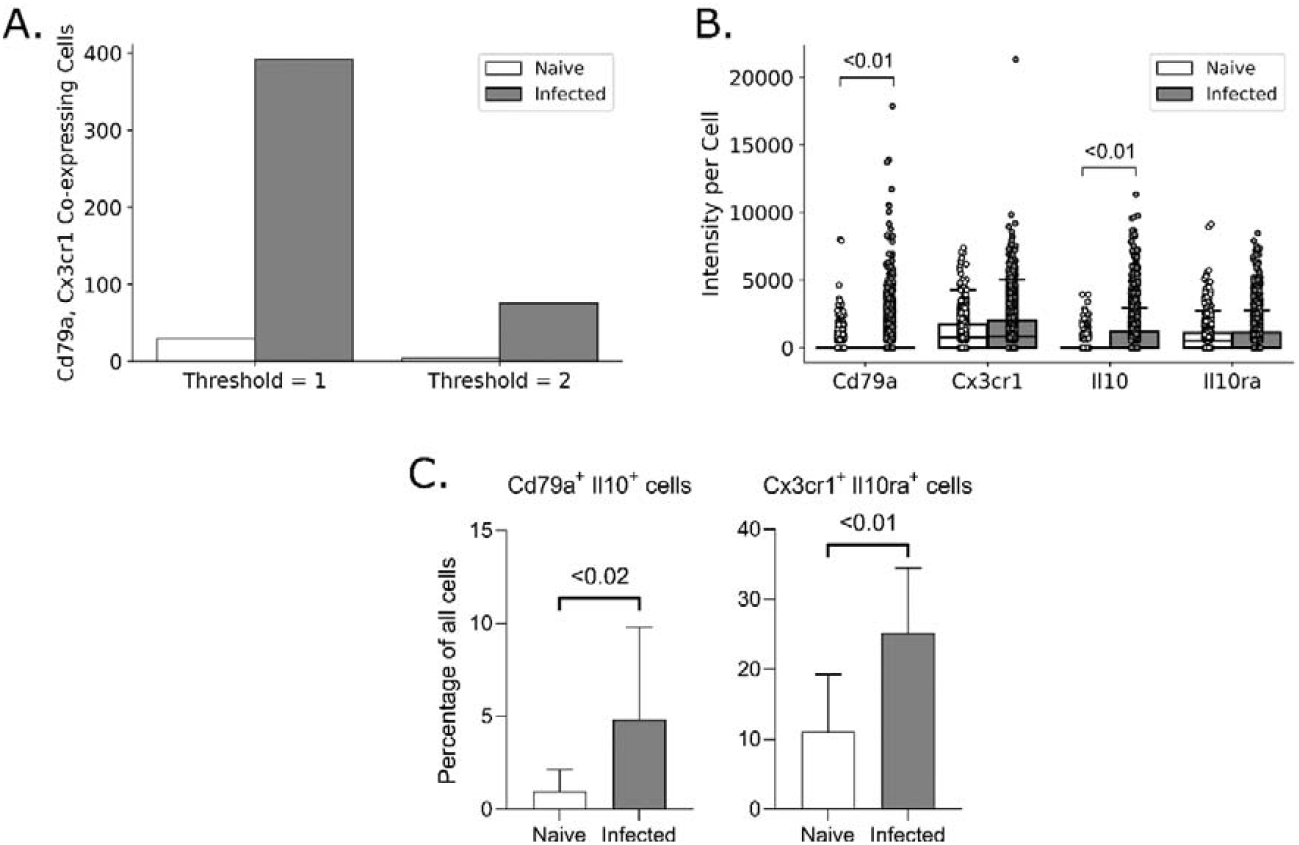
Examples of FISHtoFigure outputs from analysis of the brain dataset. **A**: Total number of cells expressing both *Cd79a* and *Cx3cr1* with threshold of either 1 or 2 transcripts per mRNA per cell. **B**: Intensities for all cells which express at least one transcript, where each point represents a cell, showing that *Cd79a* and *Il10* expression are significantly upregulated during infection (Mann-Whitney test). Box limits are defined by the interquartile range (IQR) with whiskers extending to the lowest/highest data point still within 1.5 IQR of the lower/upper quartile. This figure re-plots data originally collected in ref (4). **C**: Percentage of cells expressing both *Cd79a* and *Il10* (Bregs) and percentage expressing both *Cx3cr1* and *Il10ra* (Microglia), showing that both are significantly upregulated during infection (Mann-Whitney test). Percentages were taken from each image in the naïve and infected datasets individually and statistical analysis was performed in GraphPad PRISM.

Finally, as a demonstration of the application of FISHtoFigure in an experimental workflow, we re-analysed data that we had collected as part of a study of Breg-microglia crosstalk in the brains of mice infected with *T. b. brucei* (4). Briefly, single cell and spatial transcriptomic analyses of infected mice revealed an upregulation of the anti-inflammatory cytokine *Il10*, along with Breg and microglia associated transcripts, in the brains of *T. b. brucei* infected mice. We tested the hypothesis that during infection *Il10* expression governed crosstalk between Bregs and microglia in the brain, using smFISH and FISHtoFigure to investigate the localisation of transcripts. FISHtoFigure’s “Transcript abundance analysis” function revealed a statistically significant upregulation in *Cd79a* and *Il10* expression in infected specimens compared to naïve controls, in agreement with results from single cell transcriptomics (4). Graphical outputs in the format produced by FISHtoFigure are presented in **Figure 3B** (p<0.01, Mann-Whitney test; data from ref (4)). We then used a variety of analyses to validate that this crosstalk was driven by two specific cell types (*Il10*^+^ Bregs and *Il10ra*^+^ microglia), including visualising these cell types using smFISH. Here, we expand on this analysis by using FISHtoFigure to directly quantify the abundance of two different double-positive cell types in infected and naïve mice. We used FISHtoFigure’s “Multi-target transcript abundance” feature to analyse *Cd79a*^+^ *Il10*^+^ Breg populations and *Cx3cr1*^+^ *Il10ra*^+^ microglia populations in naïve and infected specimens. This analysis confirmed that during infection there was an upregulation of both *Cd79a*^*+*^ *Il10*^*+*^ Bregs (**Figure 3C**; p<0.02, Mann-Whitney test) and *Cx3cr1*^*+*^ *Il10ra*^*+*^ microglia (**Figure 3C**; p<0.01, Mann-Whitney test). In the context of the current paper, this demonstrates that FISHtoFigure can accurately quantify the abundance of specific cell types, including those with irregular boundaries, using multiplex expression profiles. Taken together, these findings demonstrate the value of FISHtoFigure in an experimental workflow.

## Discussion

FISHtoFigure automates the extraction and processing of transcriptomic data from QuPath-quantified smFISH data, allowing users to analyse specific transcript expression profiles in datasets that would otherwise be very difficult to parse.

Our tool is capable of analysing smFISH data by any number of mRNA targets and quantifying cell types and expression profiles with a high accuracy. Furthermore, the graphical user interface allows users to specify a positivity threshold for transcript abundance analysis (i.e., the number of transcripts required for a cell to be marked as positive, and by extension, be included in analysis), allowing users to directly control the sensitivity of the FISHtoFigure platform individually for each set of analyses.

Current analysis packages for smFISH data are largely focused on quantification and labelling of transcripts and only offer limited downstream transcript abundance analysis options, which require bioinformatics experience to implement. For example, *FISH-quant* provides a means to detect transcripts in smFISH data and assign individual transcripts to cells and subcellular compartments (11). *FISH-quant* offers downstream analysis options for mRNA expression, but this analysis is largely focused on the intracellular distributions of transcripts rather than the quantification of cells that express multiple mRNA targets. Another smFISH analysis tool, *dotdotdot*, outputs quantified cell and transcript data in a format interpretable using R or Python. However, bioinformatics experience is required to implement downstream analysis (12).

FISHtoFigure facilitates custom differential transcript and cell type abundance analyses without the need for custom code. By providing multi-transcript analysis tools in an intuitive package, FISHtoFigure significantly broadens the accessibility of smFISH analysis.

Comparison of FISHtoFigure’s spatial distribution plots with the confocal microscopy images from which they were derived demonstrates high levels of concordance between raw and quantified data (**Figure 1B**). We demonstrate that FISHtoFigure can accurately determine cell profiles in different biological systems, and that the in-built thresholding feature can substantially reduce mis-categorisation caused by the close proximity of different cell types (**Figure 2**). In the spleen dataset, applying a threshold of 2 transcripts per mRNA target per cell completely removed all mis-categorised cells in the naïve dataset and reduced mis-categorisation by >60% in the infected dataset (**Figure 2C**). In the brain dataset, applying a threshold of 2 transcripts per mRNA target per cell reduced mis-categorisation of cell types by >80% in both the naïve and infected datasets (**Figure 3A**).

In the brain dataset, FISHtoFigure enabled rapid analysis of smFISH data which would otherwise require considerable time investment and bioinformatic experience. FISHtoFigure analysis reveals a statistically significant (p < 0.01, Mann-Whitney test) upregulation in expression of *Cd79a* and *Il10* during infection (**Figure 3B**).The ability to analyse and plot cellular information for specific cell types with multiplex transcriptional profiles allowed us to identify the upregulation of *Cd79a*^*+*^ *Il10*^*+*^ Bregs and *Cx3cr1*^*+*^ *Il10ra*^*+*^ microglia in infected specimens compared with controls, a difference which would otherwise require custom code to assess (**Figure 3C**).

### Considerations and Limitations

Regarding identifying cell boundaries, QuPath has the capacity to quantify cell boundaries based on a range of factors. Here, cell nuclei were identified via fluorescent DAPI staining, and cell boundaries were approximated by applying a set radius to each identified nucleus using the “Cell Detection” function in QuPath. Though we demonstrate that this can allow the accurate quantification of cells, even for cell types with irregular boundaries, further improvements in the determination of cell boundaries, and by extension cell expression profiles, could likely be achieved through adjustments in sample preparation. For example, for challenging cell types users may wish to experiment with the use of membrane markers to further improve cell boundary quantification.

## Conclusion

The problem of balancing accessibility for non-specialist users and analytical scope is an important consideration in the development of software tools. Here, we present FISHtoFigure, an analytical platform for QuPath-quantified smFISH data capable of analysing specific cell types and multiplex transcriptomic profiles and of generating a variety of differential transcript abundance analytics for cells expressing a user-specified combination of mRNA transcripts. In the interest of accessibility for users with all levels of bioinformatic experience, we have created a simple graphical user interface and packaged FISHtoFigure as an executable program, thus allowing transcript expression analysis without interaction with raw quantified image data or custom analysis scripts. FISHtoFigure can therefore expand the in-house analysis capabilities of many research groups investigating transcriptomics via smFISH.

## Acknowledgements

We thank Colin Loney for his assistance in the acquisition of the images forming the spleen dataset used during validation. We also thank Ruaridh Wilson for providing feedback from the perspective of a computational scientist during the writing of this manuscript.

## Data Availability

All code involved in the production of the FISHtoFigure package and all analysis presented here is available on GitHub: https://github.com/Calum-Bentley-Abbot/FISHtoFigure.git

Data are available under the terms of the MIT open access licence (https://opensource.org/license/mit/).

